# Using Bifurcation Theory for Exploring Pain

**DOI:** 10.1101/757187

**Authors:** Parul Verma, Achim Kienle, Dietrich Flockerzi, Doraiswami Ramkrishna

## Abstract

Pain is a common sensation which inescapably arises due to injuries, as well as, various diseases and disorders. However, for the same intensity of disturbance arising due to the forgoing causes, the threshold for pain sensation and perception varies among individuals. Here, we present a computational approach using bifurcation theory to understand how the pain sensation threshold varies and how it can be controlled, the threshold being quantified by the electrical activity of a pain-sensing neuron. To this end, we explored the bifurcations arising from a mathematical model representing the dynamics of this neuron. Our findings indicate that the bifurcation points are sensitive to specific model parameters. This demonstrates that the pain sensation threshold can change as shown in experimental studies found in literature. Further investigation using our bifurcation approach coupled with experimental studies can facilitate rigorous understanding of pain response mechanism and provide strategies to control the pain sensation threshold.

## 1 Introduction

Nonlinear dynamics has been an integral element of the methodology of process control, an area in which Professor Georgakis, whom we are pleased to felicitate in this issue, has been engaged through most of his academic career. Traditionally, nonlinear dynamics and bifurcation theory have had many practical applications in chemical engineering involving dynamics and stability in chemically reacting systems [46], in particular, dynamics and control of polymerization reactions [45] including multiple steady states and oscillatory behavior, pattern formation on catalytic surfaces [18], multiple steady states in nonreactive [28,2] and reactive distillation processes [38], nonlinear oscillations in population balance systems including continuous crystallization [43, 20] and fluidized bed granulation [42, 49, 40], electrochemical processes including fuel cell systems [23], and bioreactors [39, 34, 55], among others. A model based analysis aims at understanding particular nonlinear phenomena, and it often builds the basis to design and control the processes so that they behave in a desired way. A constructive approach for the latter has been proposed by Marquardt and Monnigmann [36]. In the last few decades, however, the implications of nonlinear phenomena have expanded to even areas of translational significance with a potential for impact on health sciences. In this regard, as a reflection of this development, we cite, as an example, Christini et al. [6] who deliberated on the role of nonlinear dynamics in cardiac arrhythmia control. Likewise, the field of neuronal dynamics has been under active study through the noted Hodgkin-Huxley equations [25], which were developed more than 60 years ago. Our objective in this paper is to address this latter area in a direction that has high therapeutic significance for alleviating pain which invariably accompanies any form of injury and of various diseases. As pain is the body’s mechanism of protection from external danger, it must be regarded as inevitable. However, the threshold for pain can vary across individuals. In this work, we investigate how this threshold can be controlled using bifurcation theory.

The domain of pain and its level of intensity are inherently subjective in nature. The subjectivity of pain is in regard to its perception by a given individual; however, its reality is dependent on an objective external perturbation. This paper views pain as the consequence of a certain type of response by a neuron to an electrical perturbation for which there is much support in the literature [7]. The electrical perturbation is the result of an external stimulus, e.g. touch, which can excite the neuron. The response to the electrical perturbation is known to be governed by several electrical and biochemical factors. The Hodgkin-Huxley equations account for many of the electrical factors. These equations have the means to understand the generation of electrical signal in a neuron which is called an action potential. The temporal pattern of action potentials carries information that is ultimately transmitted to the brain, where it can be perceived as pain. The voltage-gated ion channels located in a neuron cell membrane play a significant role in the generation of these action potentials. This paper will examine the role of specific sodium and potassium voltage-gated ion channels as potential contributors to the generation of pain. It is our aim to promote the general exploration of pain using bifurcation theory by providing its application in this paper to a relatively simple scenario for pain mechanism.

Through our approach, we can identify parameters that can alter the pain sensation threshold. The results of our approach can be used to aid in designing experiments and subsequently exploring therapeutic strategies to control the threshold. In this paper, we will address the following: i) biological mechanism of pain sensation, ii) mathematical model representing dynamics of a pain-sensing neuron, iii) bifurcation analysis of the model equations, iv) identification of sensitive parameters in setting pain sensation threshold, and v) discussion of results with implications for a cure. This paper is intended for engineers new to computational neuroscience, the details of which can be found in books by Izhikevich, Keener, Dayan, Jaeger, Ermentrout, Schutter and Johnston [27, 33, 11, 29, 17, 51, 30].

## 2 The Neuroscience of Pain Sensation

Pain can be of three types: (i) nociceptive, (ii) neuropathic, and (iii) inflammatory. Nociceptive pain is the pain arising due to a noxious (potentially harmful) stimulus, such as touching an extremely hot or cold object. Neuropathic pain arises due to any nerve related injury, which may lead to hypersensitivity, tingling, or a burning sensation. Inflammatory pain arises due to release of inflammatory molecules (e.g. TNF*α*) as a result of internal tissue damage. Any form of sensation occurs through transmission of information across neurons in the nervous system, starting from the peripheral nervous system, reaching the central nervous system, and then transmitting back to the peripheral nervous system. Sensation is detected at the periphery by the sensory neurons, which is first converted to a chemical signal and subsequently to an electrical signal. The electrical signal is then transmitted to the spinal cord. In the case of a noxious stimulus, sensation is detected at the skin by the endings of specialized neurons. These specialized neurons are called nociceptive neurons, or nociceptors, which have receptors that can detect such a noxious stimulus, first proposed by Sherrington more than 100 years ago [53]. A nociceptor can detect noxious stimuli due to temperature, harmful chemicals such as acid, and extreme mechanical pressure. At the spinal cord, it is connected to another neuron to transmit the signal to the brain. There are a series of neuronal connections from the spinal cord to the brain. From the spinal cord, it first reaches the thalamus which relays the signal to the sensory cortex and thence to the motor cortex. From the motor cortex, the signal is sent back to the thalamus, then to the spinal cord and it finally reaches the targeted muscle cells through the motor neurons. The motor neurons control how the muscle cells will react to the noxious stimulus. If an individual touches something noxious, for example, a burning object, the following series of events will occur. The hot temperature sensation by the nociceptive neurons results in a signal to the brain and then back to the muscles at which point the individual feels a burning sensation and releases contact with the burning object. This pain sensation pathway is shown in Fig. 1. There are primarily two processes involved in pain: sensation and perception. The nerve endings at the skin sense a noxious stimulus and eventually transmit the signal to the brain. The brain decodes and perceives the stimulus as painful and accordingly generates a response to be sent back to the motor neurons. Throughout this paper, we focus on sensation, rather than perception. Details on the pain pathway can be found in multiple review papers [31, 50, 1, 54, 57].

**Figure 1:**
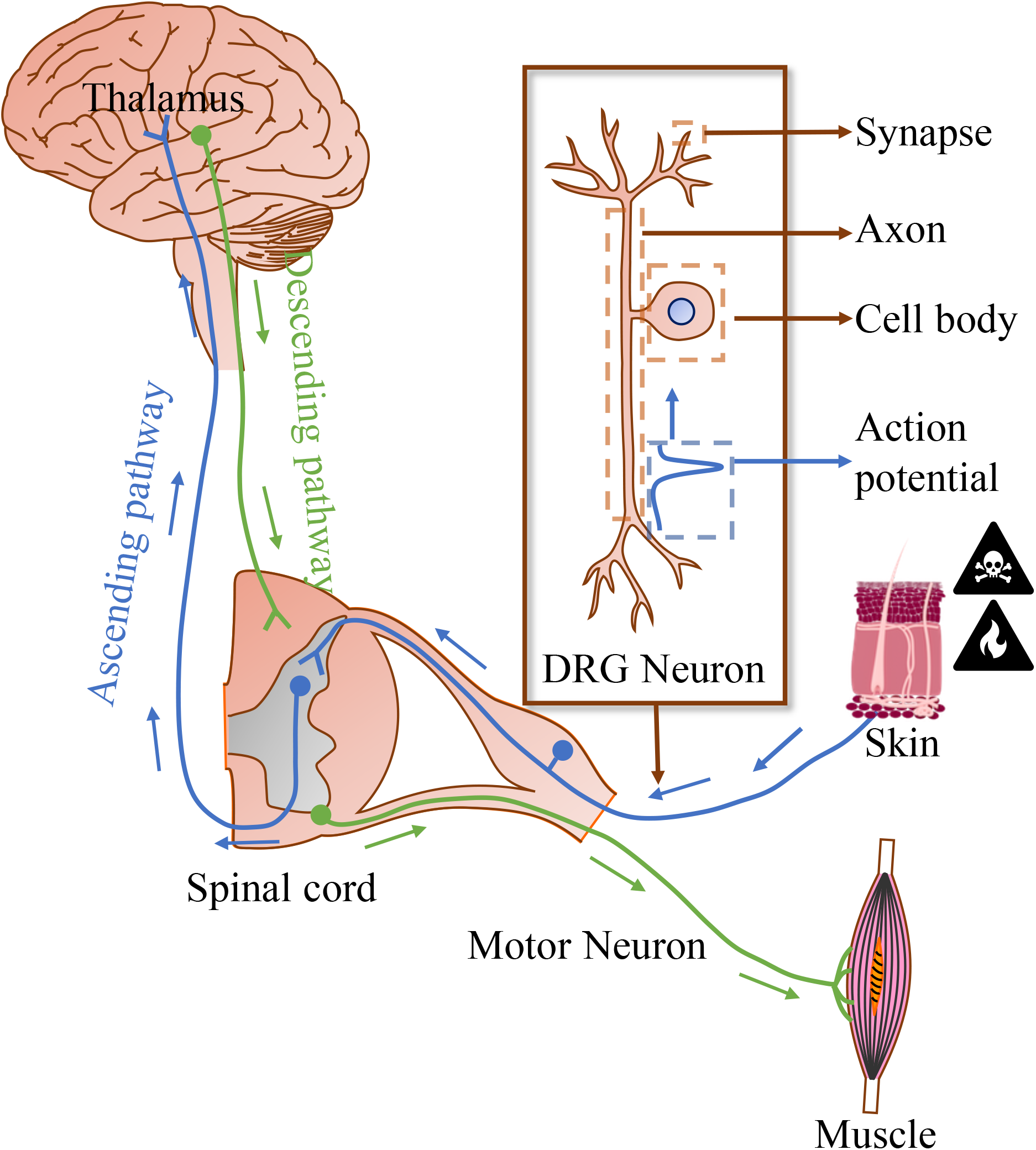
The pain sensation pathway starts from a noxious stimulus being detected by a small DRG neuron (nociceptor) at the skin. The input leads to the generation of action potentials which are then transmitted to the spinal cord. From the spinal cord, the input is further transmitted to the thalamus via the ascending pathway. The response is transmitted back to the spinal cord via the descending pathway. It is finally transmitted to the muscle cells via the motor neuron. The muscle cells respond in concordance with the signal. The DRG neuron consists of a cell body, an axon across which the action potential is transmitted, and ends at the synapse which transmits information from one neuron to the other.

Most small dorsal root ganglia (DRG) neurons are nociceptors [32], shown in Fig. 1. In this work, we will only focus on small DRG neurons. Specific alterations or injury in this neuron can lead to change in the nociceptive pain threshold (gain or loss of pain sensation), neuropathic pain, inflammatory pain, or a combination of these. These neurons consist of a cell body, an axon, and synapses at the branch endings. Furthermore, small DRG neurons are pseudo-unipolar, which means that that they have two axons branching out from the main cell body, one reaching the periphery and the other reaching the central nervous system.

Neuron signal transmission occurs when it becomes excitable due to an adequate electrical perturbation and consequently generates action potentials. An action potential is a brief peak in potential dynamics observed across a neuronal membrane. The temporal pattern (consisting of frequency and duration) and source of action potential generation determine the message being transmitted. A schematic of an action potential is shown in Fig. 2. Action potential transmission occurs within a neuron across the length of the axon and subsequently in between two neurons via their synapses. The action potential is generated due to opening and closing of voltage-gated ion channels on a neuronal membrane. The neuronal membrane consists of a lipid bilayer, and the voltage-gated ion channels are transmembrane proteins with pores that are selectively permeable to specific ions (e.g. sodium, potassium, calcium, or chloride). These ions have their respective equilibrium potentials. Sodium has a positive while potassium has a negative equilibrium potential. The resting membrane potential of a neuron is negative. These channels are shown in Fig. 3.

**Figure 2:**
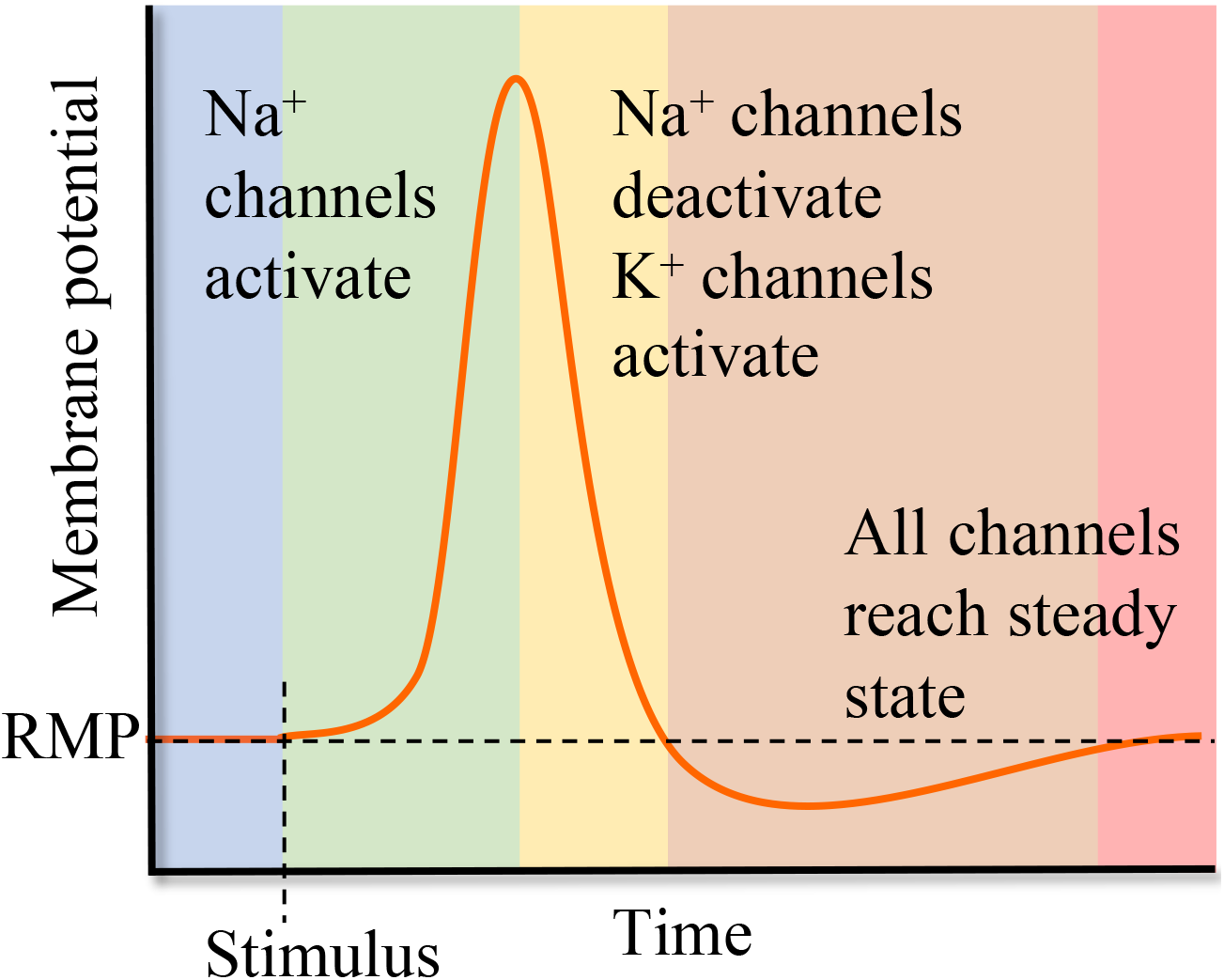
A schematic of an action potential. When a stimulus is applied, an action potential is generated due to activation of sodium channels leading to the rise in membrane potential. Following this rise, sodium channels inactivate and potassium channels activate, leading to a decrease in the potential. Finally, all channels attain steady states and the membrane reverts back to the resting membrane potential (RMP).

**Figure 3:**
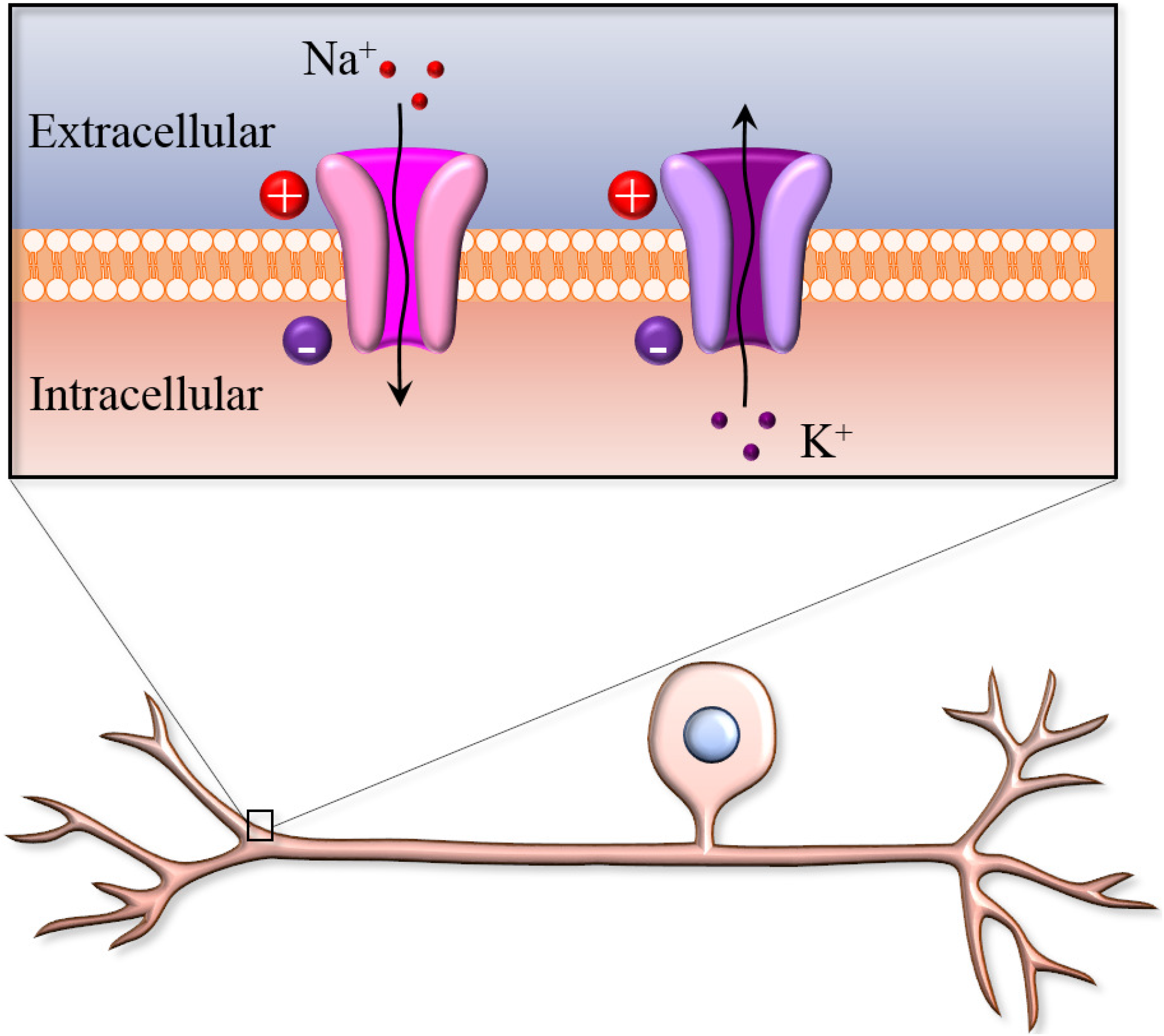
A schematic of a neuronal membrane. The membrane consists of a lipid bilayer. The voltage-gated sodium and potassium channels are transmembrane pores. The extracellular concentration of sodium is greater, leading to an inflow of sodium ions when the channel opens. The intracellular concentration of potassium is greater, leading to an outflow of potassium ions when the channel opens.

When an external stimulus is applied, sodium channels get activated and open up. This leads to a flux of sodium ions across the membrane into the neuron, raising the membrane potential towards the positive equilibrium potential of sodium. This inflow continues until it reaches a threshold which corresponds to a peak in the action potential. While reaching the peak, sodium channels start to inactivate by closing while potassium channels activate and open. Activation of potassium channels lead to an inflow of potassium ions, lowering the membrane potential towards the negative equilibrium potential of potassium. Subsequently, most of the channels close, reaching a steady state and bringing the membrane potential back to the resting membrane potential. These series of events correspond to the generation of one action potential. The timing associated with opening or closing of these channels is a function of time constants of activation and inactivation. Sodium activation has a smaller time constant and as a result activates first. Sodium inactivation and potassium activation have relatively larger time constants resulting in a time lag. This interplay of fast and slow dynamics leads to the generation of an action potential.

A stimulus can either be an interim, pulse-like disturbance in which case the membrane potential will return to the same rest point, as shown in Fig. 2. Or, it can be a permanent disturbance in which case the membrane potential will return to a new rest point which is discussed in the next section. In both the cases, action potentials are only fired if the stimulus exceeds a threshold value.

Hodgkin and Huxley were the first researchers to develop a mathematical model describing action potential generation via interplay of one sodium and one potassium channel. They developed this model for a squid giant axon. Since then, Hodgkin-Huxley type models have been developed for several other neurons, incorporating multiple channels. In this paper, we focus on a Hodgkin-Huxley type model representing dynamics of a pain-sensing neuron.

## 3 Mathematical Model of a Pain-Sensing Neuron

Here, we describe a model representing dynamics of a small DRG neuron. As previously mentioned, the model equations are of Hodgkin-Huxley type. There are multiple types of ion channels present in the human nervous system. For every ion type, a variation of channel subtypes exist, each having different kinetic properties. For this model, we included two sodium channels: Na_v_1.7 and Na_v_1.8, and two potassium channels: delayed-rectifier potassium channel (K channel) and A-type transient potassium channel (KA channel). Na_v_1.7 and Na_v_1.8 are simply two types of sodium channels with different kinetic properties, but both consist of fast activating and slow inactivating gating variables. The K channel is a Hodgkin-Huxley type potassium channel with only one activating gate variable which is activated later than the sodium channels. The KA channel has both activating and inactivating gating variables, similar to sodium channels. We further included a non-gated leak channel, as was done for the original Hodgkin-Huxley models. These specific channels were chosen since they are most prominent in a small DRG neuron membrane, and, therefore, models consisting only of these channels have been developed and analyzed previously [52, 4].

The aforementioned components of the neuronal membrane can be represented as an electrical circuit, as shown in Fig. 4. We assume the membrane to have a specific capacitance *c* (= *C*/*A*, *C* being the membrane capacitance and A the membrane surface area) with some potential *V* (= *V_in_* – *V_out_*) across it. Traditionally, the extracellular potential is assigned as zero which then defines the intracellular resting potential as negative. *I_ext_*(*t*), where *t* is time, is the external stimulus current. The direction of this current is opposite to the direction of current due to the ion channels. As shown in the figure, the sodium and potassium channels have variable conductance, while the leak channel has a constant conductance, implying it is non-gated. The equilibrium potential of the leak channel is negative as well, in order to bring the membrane potential back to the negative resting membrane potential. The variable conductance are functions of the membrane potential, described by Hodgkin-Huxley type equations.

**Figure 4:**
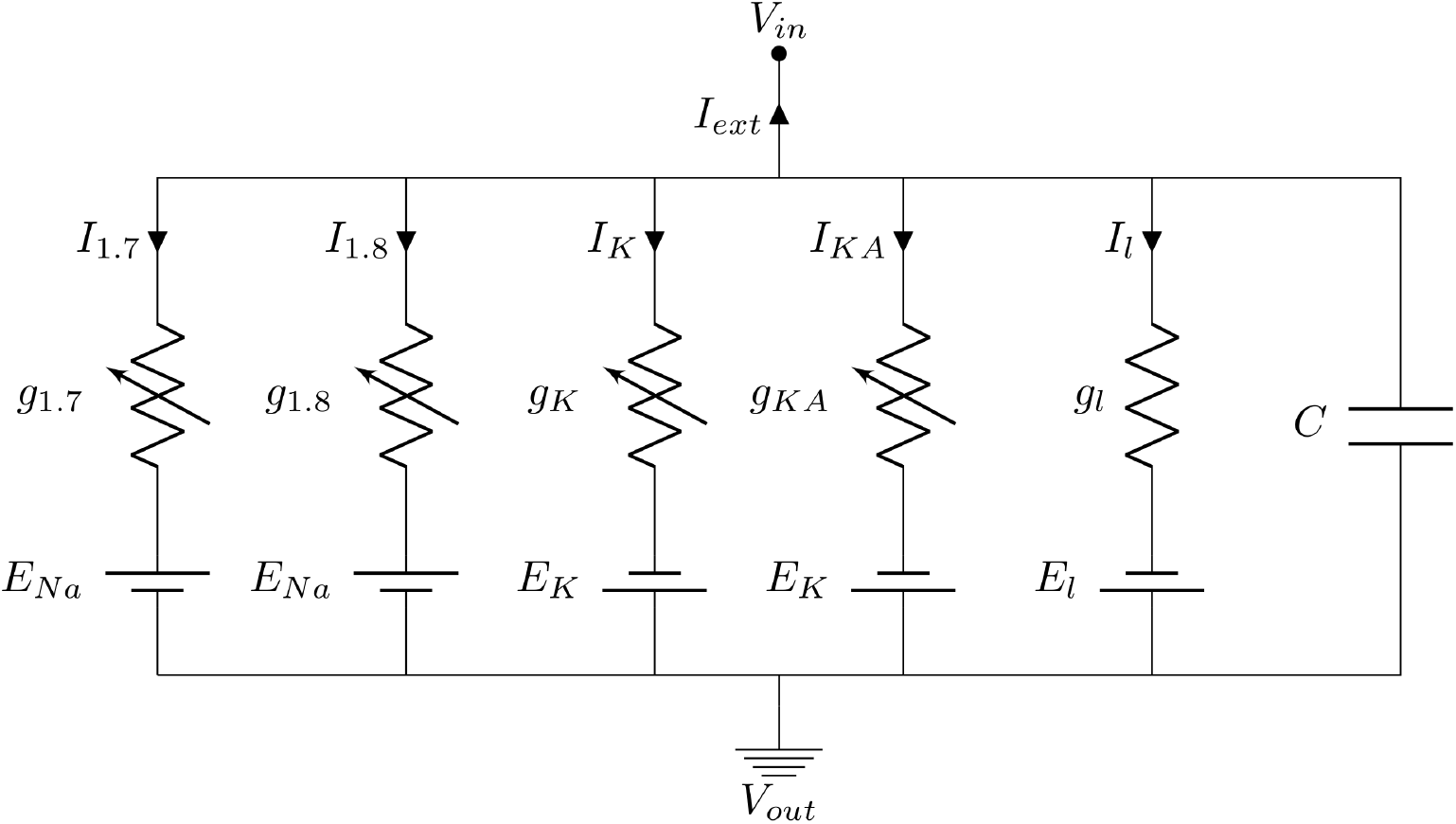
Circuit diagram representing neuron membrane. *V_in_*: intracellular potential; *V_out_*: extracellular potential; *I*_1.7_, *I*_1.8_, *I_K_*, *I_KA_*, *I_l_*: current due to Na_v_1.7, Na_v_1.8, delayed rectifier potassium, A-type transient potassium and leak channels respectively; *I_ext_*: external stimulus current; *g*_1.7_, *g*_1.8_, *g_K_*, *g_KA_*, *g_l_*: conductance of Na_v_1.7, Na_v_1.8, delayed rectifier potassium, A-type transient potassium and leak channels respectively; *E_Na_*, *E_K_*, *E_l_*: equilibrium sodium, potassium and leak potentials respectively, *C*: membrane capacitance.

Using Kirchhoff’s law, the equation for potential dynamics can be written as the following:

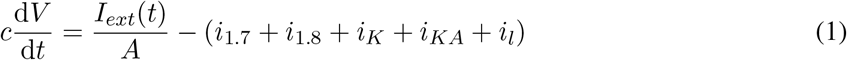

The individual specific ionic currents are defined as following:

1. 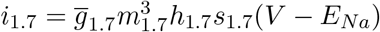
2. 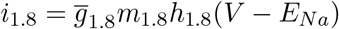
3. 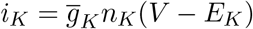
4. 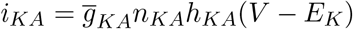
5. 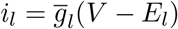

Here, 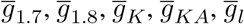 are the maximal specific conductance, taken as constants. The gating variables *m*_1.7_, *h*_1.7_, *s*_1.7_, *m*_18_, *h*_1.8_, *n_K_*, *n_KA_*, *h_KA_* lie between 0 and 1, and are functions of *V*. *E_Na_*, *E_K_* and *E_l_* are ion equilibrium potentials. The final equation for membrane potential is the following:

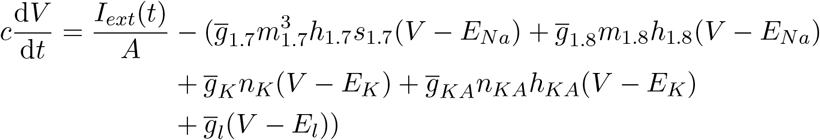

The gating variables correspond to fractions of ion channels that are open. Suppose *x* is a gating variable, where *x* = *m*_1.7_, *h*_1.7_, *s*_1.7_, *m*_1.8_, *h_1.8_*, *n_K_*, *n_KA_*, *h_KA_*. Then, the equation for *x* is written as:

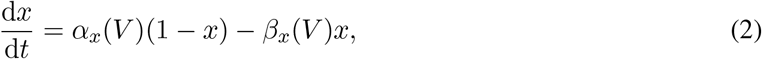

where, *α_x_* (*V*) is the rate of opening of gate *x* and *β_x_* (*V*) is the rate of closing. On rearranging this equation, we get the following:

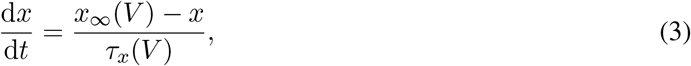

where, 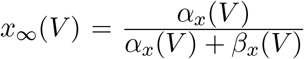 and 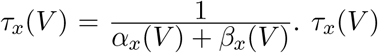 is the potential-dependent time constant. Typically, *α_x_* (*V*) and *β_x_* (*V*) are assumed to be of the following form:

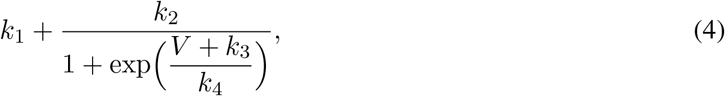

where, for each *x*, the corresponding *k*_1_ to *k*_4_ are constants. In total, this model consists of 9 ordinary differential equations.

When *I_ext_* = 0, the resting membrane potential of this system will be the following, by setting all the derivatives to zero:

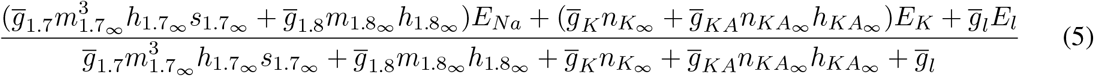

The equations for gating variables and the parameter values are mentioned in the supplementary document, and have been obtained from previous models [52, 4, 48]. Typically, channel kinetics (values of *k*_1_, *k*_2_, *k*_3_, *k*_4_) are obtained from voltage-clamp experiments. In these experiments, the potential across a neuron is fixed and then it is stepped to a different value, and the corresponding current pattern evolution is observed. This data is fit to the model equations to determine the constants. We do not describe the details of voltage-clamp experiments and subsequent parameter estimation here, but recommend the seminal paper by Hodgkin and Huxley for details on how the original equations were obtained [25], and the book by Johnston and Wu for more details on the neurophysiology [30]. Throughout this paper, the units of potential are mV, specific capacitance mS/cm^2^, current pA, and time ms.

*m*_1.7_ and *m*_1.8_ are fast activation variables of sodium, *h*_1.7_ and *h*_1.8_ are slow inactivation variables of sodium, and *s*_1.7_ is ultra-slow inactivation of sodium 1.7 channel. *n_K_* and *n_KA_* are activation variables of potassium channels and *h_KA_* is inactivation variable of KA channel. The gating variables *n_K_* and *n_KA_* are slower than those of sodium channels, which ensures the descent after the peak in the action potential. We discuss this below by comparing Fig. 5C-F and by evaluating approximate time scale of evolution of these variables in the supplementary document.

**Figure 5:**
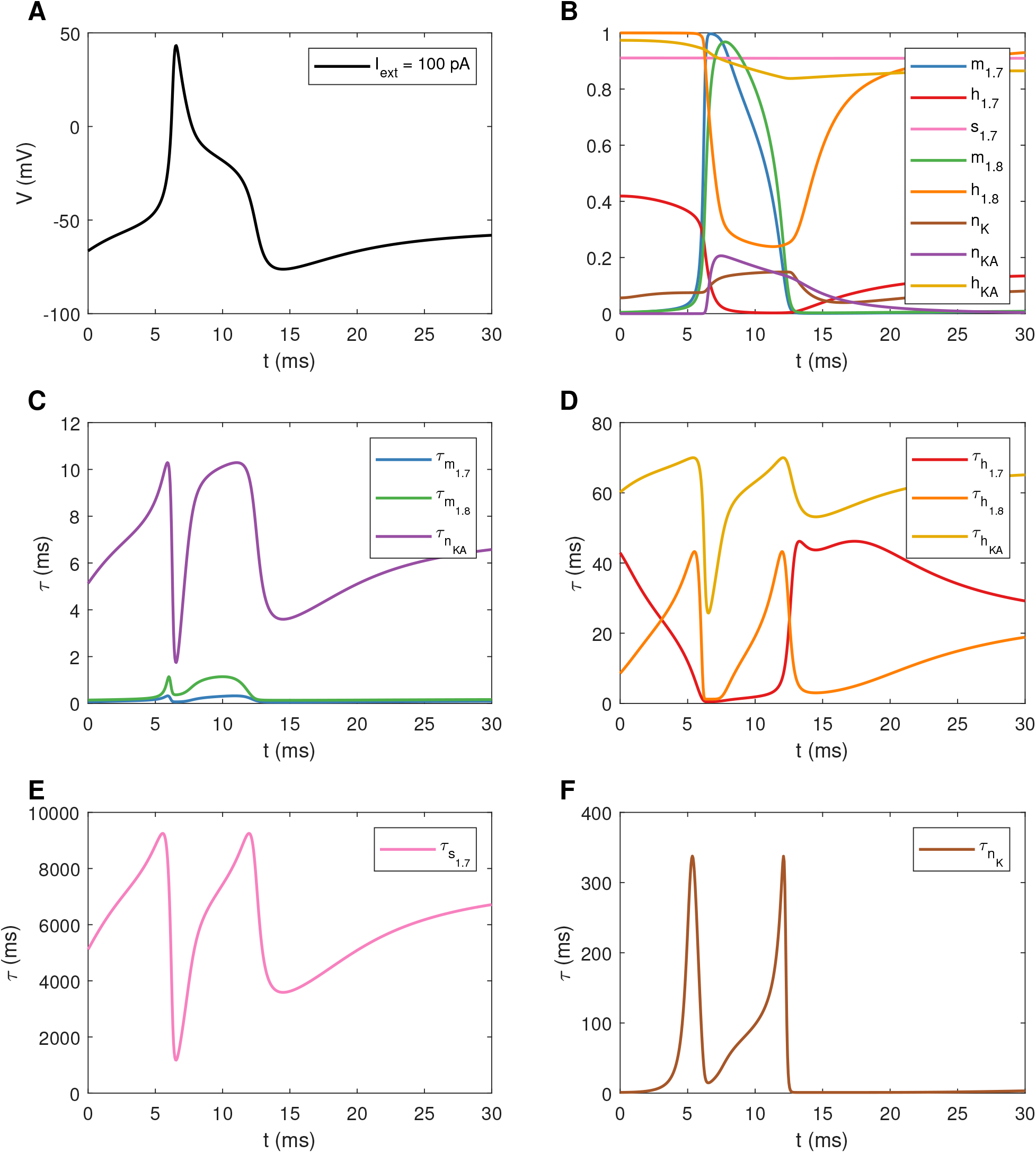
A: Action potential generated due to a constant external current *I_ext_* = 100 pA, B: Dynamics of activation and inactivation state variables, C-F: Dynamics of potential-dependent time constants for the state variables.

Upon simulating the system with a constant external current source of 100 pA (starting from *t* = 0 ms), an action potential is generated, shown in Fig. 5A. The initial conditions here correspond to those for *I_ext_* = 0. At around 5 ms, membrane potential begins to shoot up drastically. At this time, as shown in Fig. 5B, the state variables *m*_1.7_ and *m*_1.8_ begin to increase. Simultaneously, *h*_1.7_ and *h*_1.8_ start decreasing and remain low in the region between around 7-12 ms, corresponding to the relatively flat region in the descent of the action potential. During the descent, both *n_K_* and *n_KA_* increase, corresponding to opening of potassium channels. The temporal pattern of *h_KA_* is similar to *h*_1.7_ and *h*_1.8_. *s*_1.7_ remains relatively constant in this time frame. Eventually, these state variables reach steady state values, and the updated resting membrane potential, for a given external input (in the form of a step function), is the following:

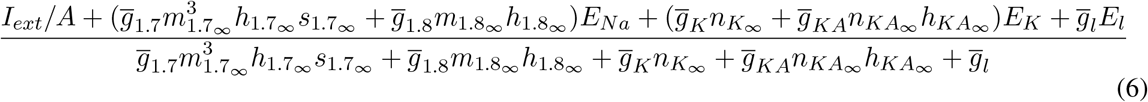

The temporal pattern of the potential-dependent time constants are shown in Fig. 5C-F. *m*_1.7_ and *m*_1.8_ have relatively smaller time constants, implying faster kinetics as seen in Fig. 5B, ensuring activation of sodium channels first. The remaining time constants for *n_KA_*, *h_KA_*, *h*_1.7_, *h*_1.8_ are larger, implying slow activation of the KA channel and slower inactivation of KA and both the sodium channels. The K channel inactivates even slower, as seen in Fig. 5F by the magnitude of its time constant. *s*_1.7_ is the slowest with time constant larger than all other variables, leading to ultra-slow inactivation of Na_v_1.7 channel. Therefore, *s*_1.7_ remains relatively constant as shown in Fig. 5B.

Periodic firing of action potentials by this neuron is associated with pain of some degree. In order to understand the onset of periodic firing, we applied bifurcation theory to this model, using *I_ext_*, *E_Na_* and *E_K_* as the primary bifurcation parameters. In this section, we will demonstrate some dynamic simulations to show how the system dynamics vary upon changing these parameters. First, we varied *I_ext_*, assuming it to be a time-independent step function. As shown in Fig. 6A, for a small value of *I_ext_*, one action potential is produced after which the system approaches a steady state. Upon increasing *I_ext_*, the system displayed various types of mixed-mode oscillations (MMO) consisting of small amplitude oscillations around the lowest unstable steady state solution (see Fig. 9) and full-blown action potentials, as shown in Fig. 6B. For specific values of *I_ext_*, we have also observed more complicated attractors which we will not investigate here. Upon further increasing *I_ext_*, the system displayed periodic firing of action potentials (Fig. 6C), indicative of pain of higher intensity as compared to that in Fig. 6B. In general, dynamical simulations suggest that the average frequency of the large amplitude oscillations is increasing with *I_ext_* (corresponding to increase in pain intensity). The bottom sub-figures show phase portraits with three state variables: *V*, *n_K_* and *h*_KA_. For 6A, since there are no oscillations, a closed curve is not formed in the phase portrait. The phase portrait in 6B consists of a closed curve with extra smaller loops to take into account the small amplitude oscillations. The phase portrait in 6C is a single closed curve corresponding to the large amplitude oscillations of action potentials.

**Figure 6:**
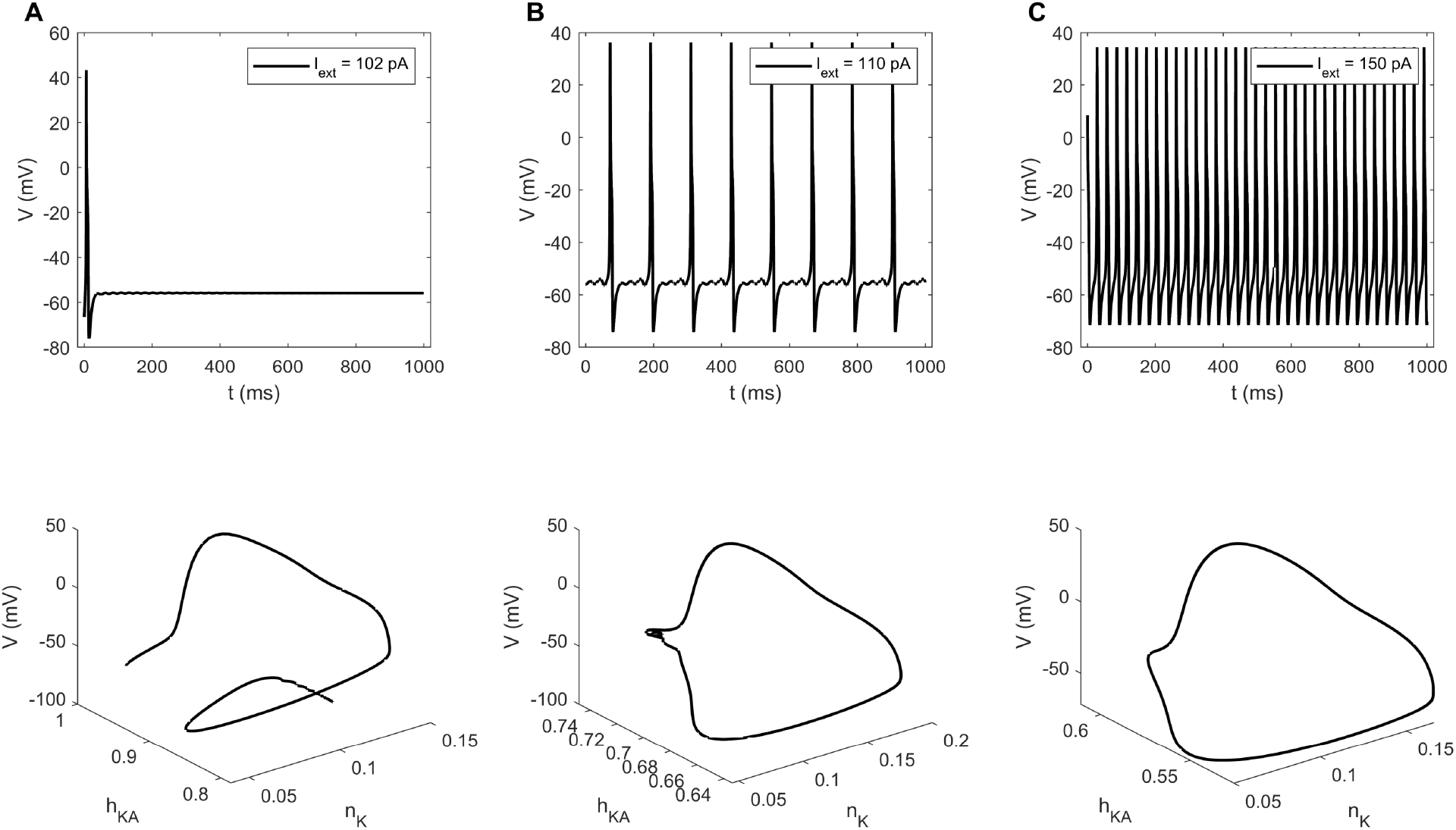
Dynamic simulations for A: *I_ext_* = 102 pA, B: *I_ext_* = 110 pA, C: *I_ext_* = 150 pA

Similar dynamical patterns are observed upon varying *E_Na_* and *E_K_*. For smaller values of *E_Na_*, a single action potential is generated, as shown in Fig. 7A. Increasing *E_Na_* leads to MMO and then periodic firing, as shown in Fig. 7B and Fig. 7C respectively. A similar pattern is observed in the case of *E_K_*, as shown in Fig. 8. The phase portraits look similar as well.

**Figure 7:**
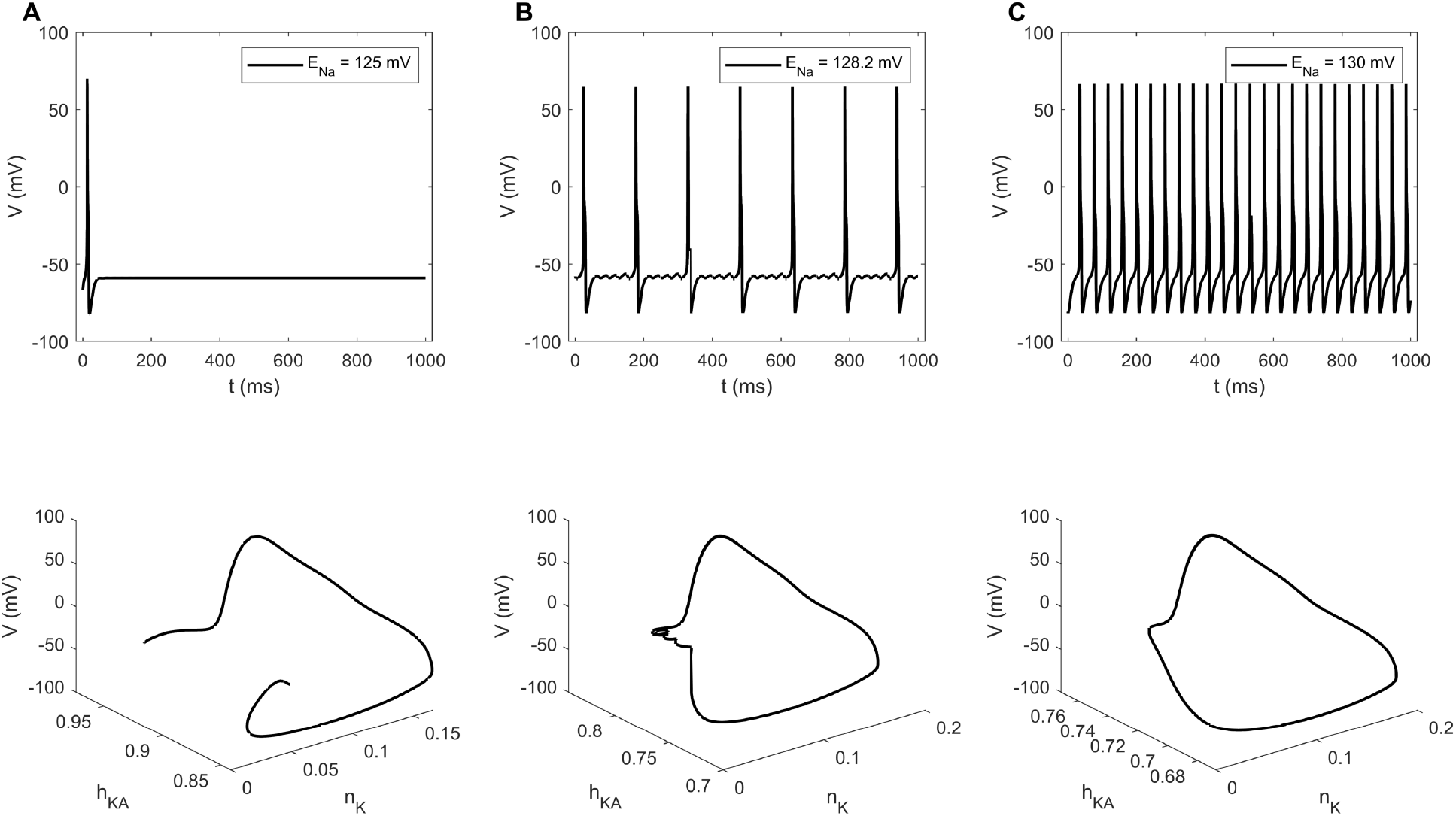
Dynamic simulations for A: *E_Na_* = 125 mV, B: *E_Na_* = 128.2 mV, C: *E_Na_* = 130 mV

**Figure 8:**
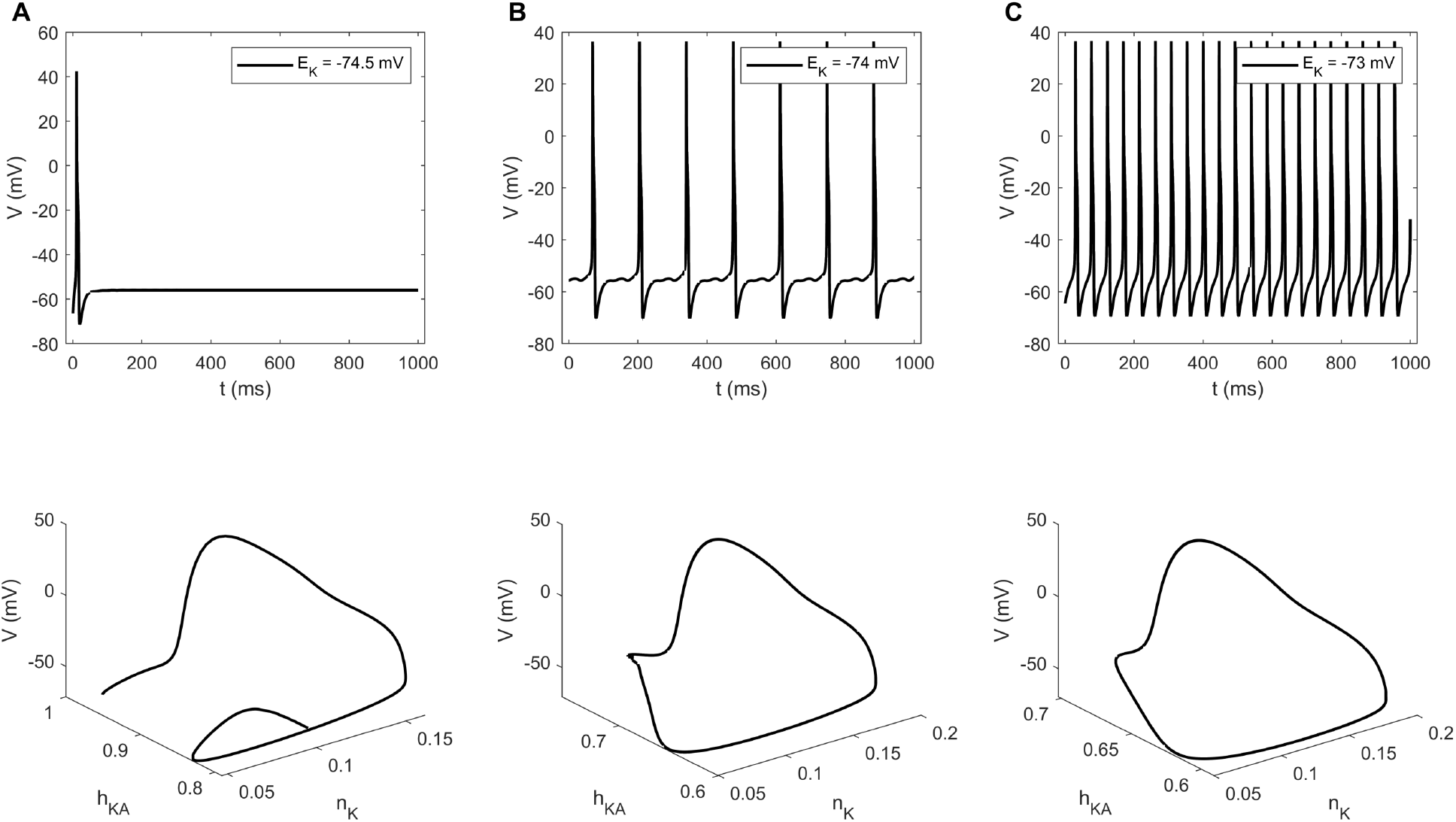
Dynamic simulations for A: *E_K_* = −74.5 mV, B: *E_K_* = −74 mV, C: *E_K_* = −73 mV

## 4 Influence of *I_ext_*

We first performed a one-parameter continuation of solutions with *I_ext_* as the primary bifurcation parameter, using XPPAUT [16]. *I_ext_* is assumed to be time independent. The partial bifurcation diagram is shown in Fig. 9. First, a continuation of steady state solutions was done starting from a stable steady state at the left of Fig. 9. The red branch of stable steady state solutions loses its stability at a subcritical Hopf bifurcation point (HB in Fig. 9 at around *I_ext_* = 103 pA) and undergoes subsequent hysteresis with two limit points (LP in Fig. 9) leading to a multiplicity of unstable steady state solutions in between (in black). At the HB point, a branch of stable steady states coalesces with a branch of unstable period solutions (in blue), with the blue branch undergoing a cyclic limit point (CLP in Fig. 9) at its turning point. Starting at the HB point, the blue periodic branch has two unstable directions, and beyond the CLP, it has either one or three unstable directions due to one real characteristic value passing through zero. At the end of this branch, the period increases massively. It is, therefore, conjectured that this branch ends in a period-infinity solution.

**Figure 9:**
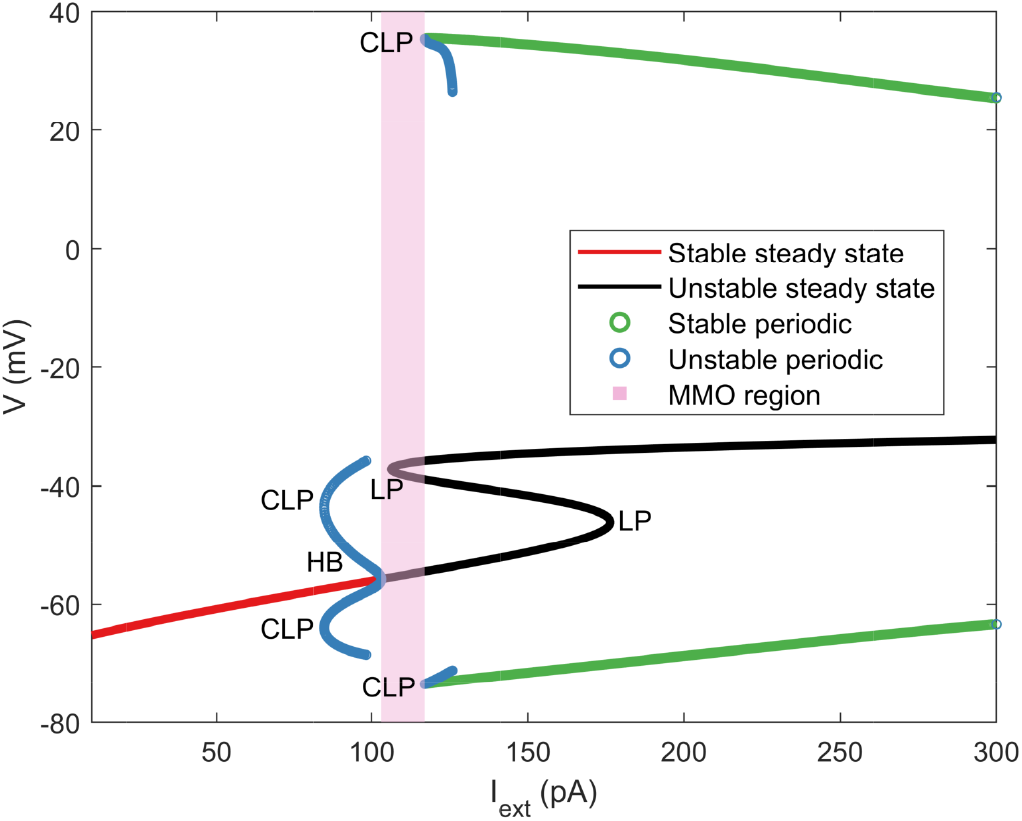
Partial bifurcation diagram with *I_ext_* as the bifurcation parameter. HB: Subcritical Hopf bifurcation point (at *I_ext_* = 102.99 pA), LP: Limit point, CLP: Cyclic limit point (at *I_ext_* = 116.98 pA for the stable periodic branch).

Furthermore, there is a branch of stable large amplitude oscillations indicated by the green circles in Fig. 9, which corresponds to the periodic firing of action potentials as illustrated in Fig. 6C. The branch of large amplitude oscillations becomes unstable at a CLP (Fig. 9 at around *I_ext_* = 117 pA) and ends shortly after a period-infinity solution. Between the HB point and this CLP, MMO (Fig. 6B) are observed as well as more complicated patterns of behavior associated with these types of MMO. Bifurcations and solution branches corresponding to the MMO region are currently under investigation and are, therefore, not shown in Fig. 9.

In this article, we focus on how these detected bifurcation points are helpful in understanding the pain sensation. The bifurcation points separate the “no-pain” (stable steady state) region from the “painful” (oscillation of action potentials) region, where the intensity of pain increases upon increasing *I_ext_* since it is determined by the frequency of firing of action potentials. Hence, understanding how these bifurcation points can shift can indicate how the pain threshold changes, due to factors such as external injury or genetic mutations. In order to explore this, we focus on the following bifurcation points: HB point, LP on the unstable steady state branch, and CLP of the stable periodic branch. We performed two-parameter continuations of these points to demonstrate how they shift on changing specific model parameters, possibly shifting the pain sensation threshold.

We further focus on finding specific kinetic parameters in sodium channels that can shift the bifurcation points. This is because several mutations in sodium channels have been found to be associated with loss or gain of pain sensation [3, 15, 10]. Most of the previous work in this field is based on experiments and computational modeling. There is limited literature on use of bifurcation theory to understand the role of pain sensation [47, 44, 58]. Here, we present an approach using bifurcation theory to examine some of these mutations.

### 4.1 Na_v_1.7 Channel Mutations

Na_v_1.7 is a widely studied channel and a myriad of mutations in Na_v_1.7 have been found to be associated with gain or loss of pain sensation [14]. Gain of pain sensation implies that the pain sensation threshold is relatively lower, while individuals with loss of pain sensation might not feel any pain even after severe injury. Such a mutation was identified in specific individuals who had never experienced pain [8]. The gain of pain sensation case is associated with discomfort even without being in a dangerous scenario. The loss of pain sensation is associated with multiple severe injuries with little to no reaction. In both cases, the pain sensation threshold no longer meets the physiological requirements. In order to understand how the pain threshold can change due to mutations in Na_v_1.7, we varied specific parameters in the activation variable *m*_1.7_. The kinetics of *m*_1.7_ is described by *α*_*m*_1.7__ (*V*) and *β*_*m*_1.7__ (*V*), which are written in the form of Eq. 4. To investigate mutation in kinetics of *m*_1.7_, we varied *k*_3_ by introducing a dummy variable *v*_0_, first in the equation for *α*_*m*_1.7__ (*V*), such that it becomes:

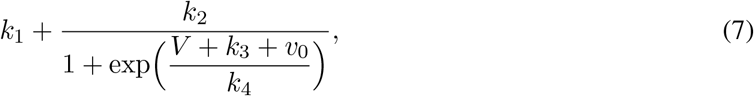

We then performed a two-parameter continuation of bifurcation points with *v*_0_ and *I_ext_*, shown in Fig. 10A. Similarly, we investigated mutation in *β*_*m*_1.7__ (*V*) kinetics, by introducing the dummy variable exclusively in its kinetics. We again performed two-parameter continuation with *v*_0_ and *I_ext_*, shown in Fig. 10B. For both the cases, it can be seen that the bifurcation points shift to the left upon increasing *v*_0_. This leads to a lower bifurcation point, implying the neuron will start periodic firing of action potentials with a lesser stimulus. This would lead to a decrease in the threshold for pain sensation.

**Figure 10:**
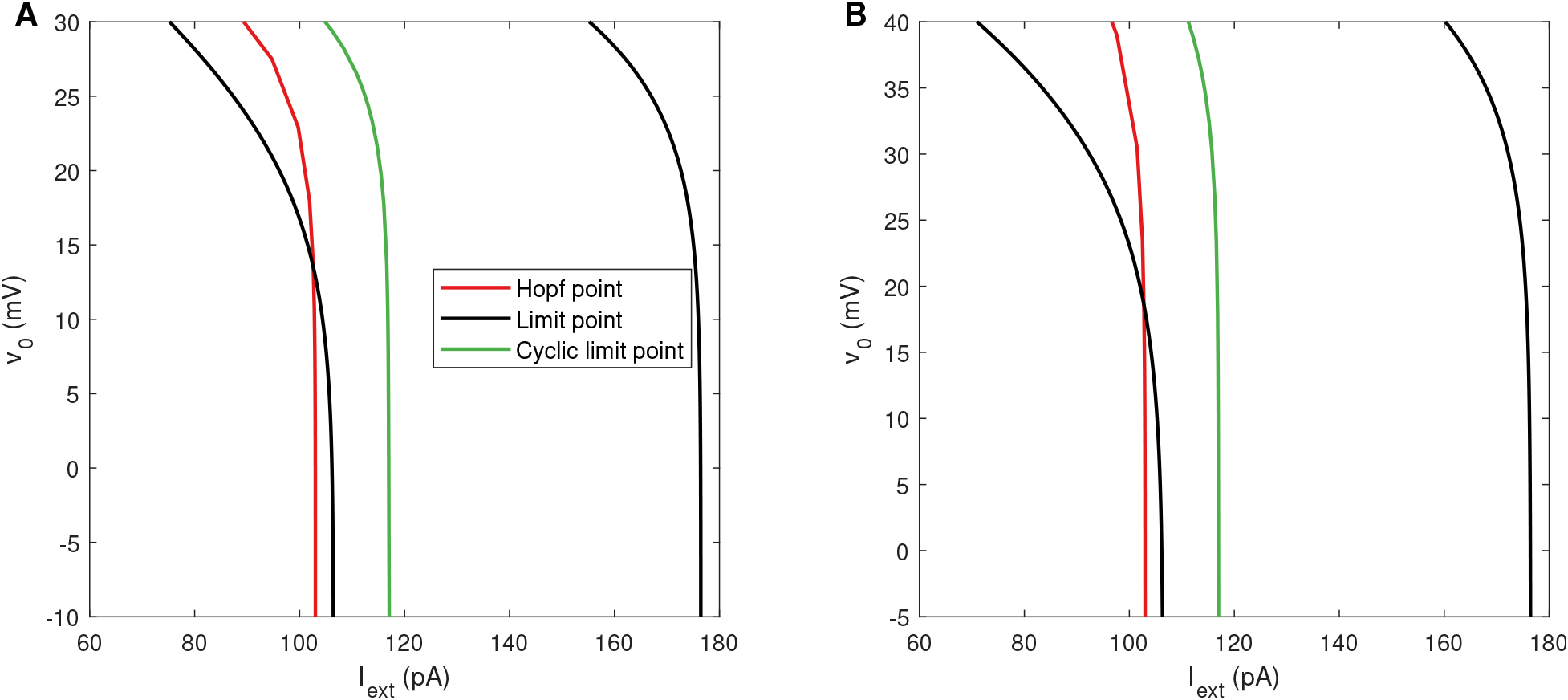
Two parameter continuation for *v*_0_ in A: *α*_*m*_1.7__ (*V*) and B: *β*_*m*_1.7__ (*V*). The plot shows how the bifurcation points vary upon changing *v*_0_.

Increasing *v*_0_ shifts the steady state activation variable 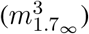 versus potential plot to the left, as shown in Fig. 11. Fig. 11 is a plot of activation variable versus potential during the ascent of the action potential. Association between the leftward shift of the steady state activation variable and the gain of pain sensation has been seen in several experiments, where mutations in Na_v_1.7 led to chronic pain associated with a burning sensation in individuals [9, 13, 5, 24, 21, 52, 56]. Thus, these results are in concordance with existing experimental findings.

**Figure 11:**
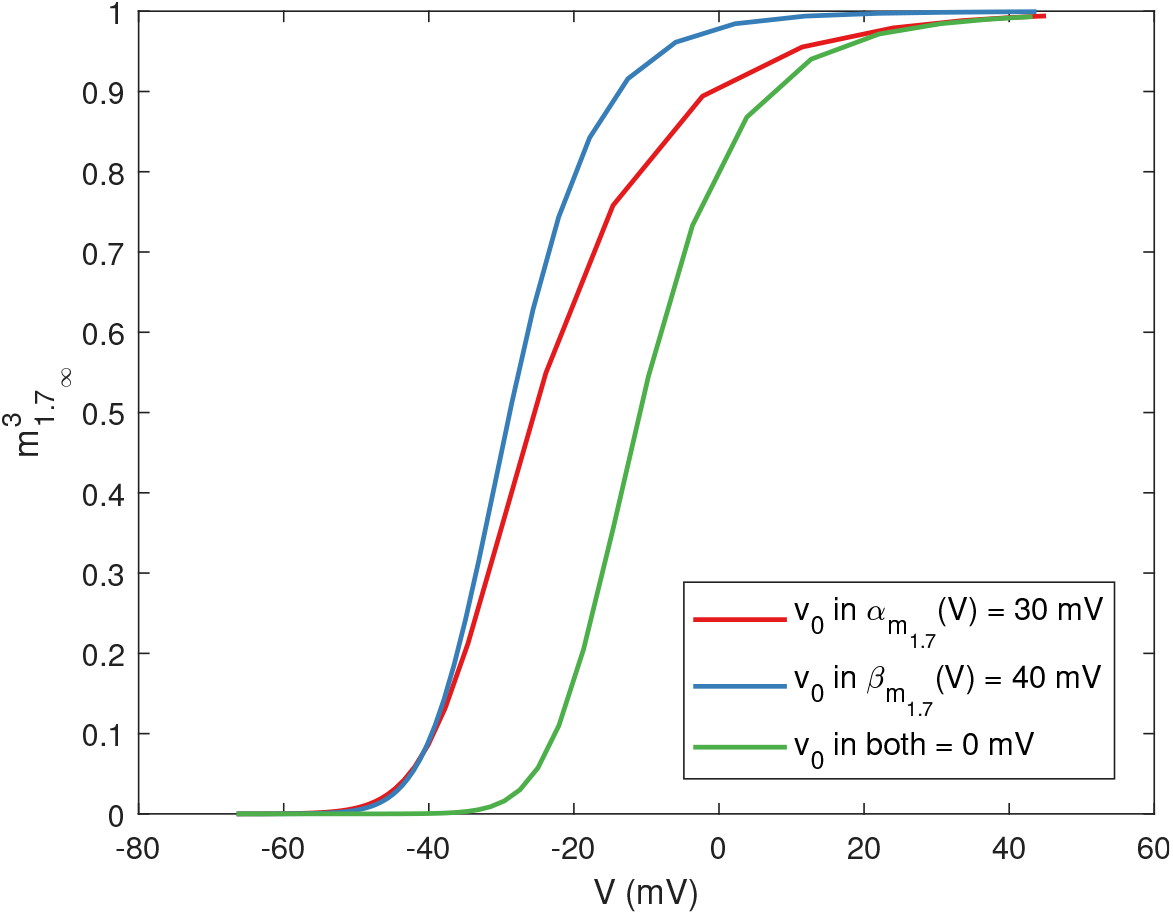
There is a shift in 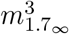 as a result of an increase in *v*_0_.

### 4.2 Na_v_1.8 Channel Mutations

Na_v_1.8 channel has also been associated with alterations in pain sensation [22]. However, it is not as well studied as Na_v_1.7. Na_v_1.8 is associated primarily with inflammatory pain. Here, we similarly introduced dummy variable *v*_0_ one by one in both *α*_*m*_1.8__ (*V*) and *β*_*m*_1.8__ (*V*). As shown in Fig. 12, the bifurcation points shift to the left, indicative of a decrease in pain sensation threshold. Furthermore, for lower values of *v*_0_, the CLP is to the left of the HB point (see Fig. 12), which indicates bistability.

**Figure 12:**
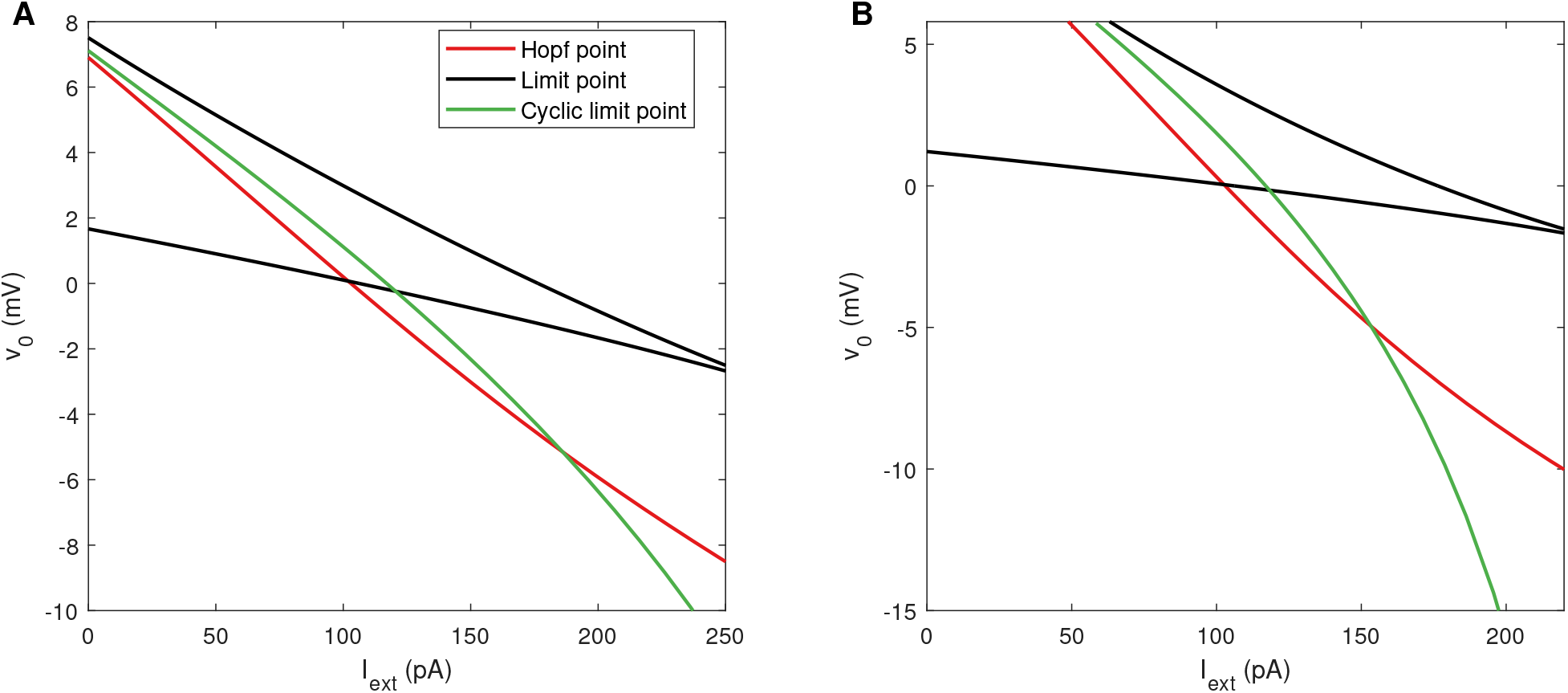
Two parameter continuation for *v*_0_ in A: *α*_*m*_1.8__(*V*) and B: *β*_*m*_1.8__(*V*). The plot shows how the bifurcation points vary upon changing *v*_0_.

Similar to the Na_v_1.7 analysis, a leftward shift in the plot of steady state activation variable *m*_1.8_∞__ versus membrane potential can be seen in Fig. 13. Again, such a shift has been reported in literature, corresponding to a gain of sensation mutation in Na_v_1.8, associated with painful neuropathy in individuals [19, 26].

**Figure 13:**
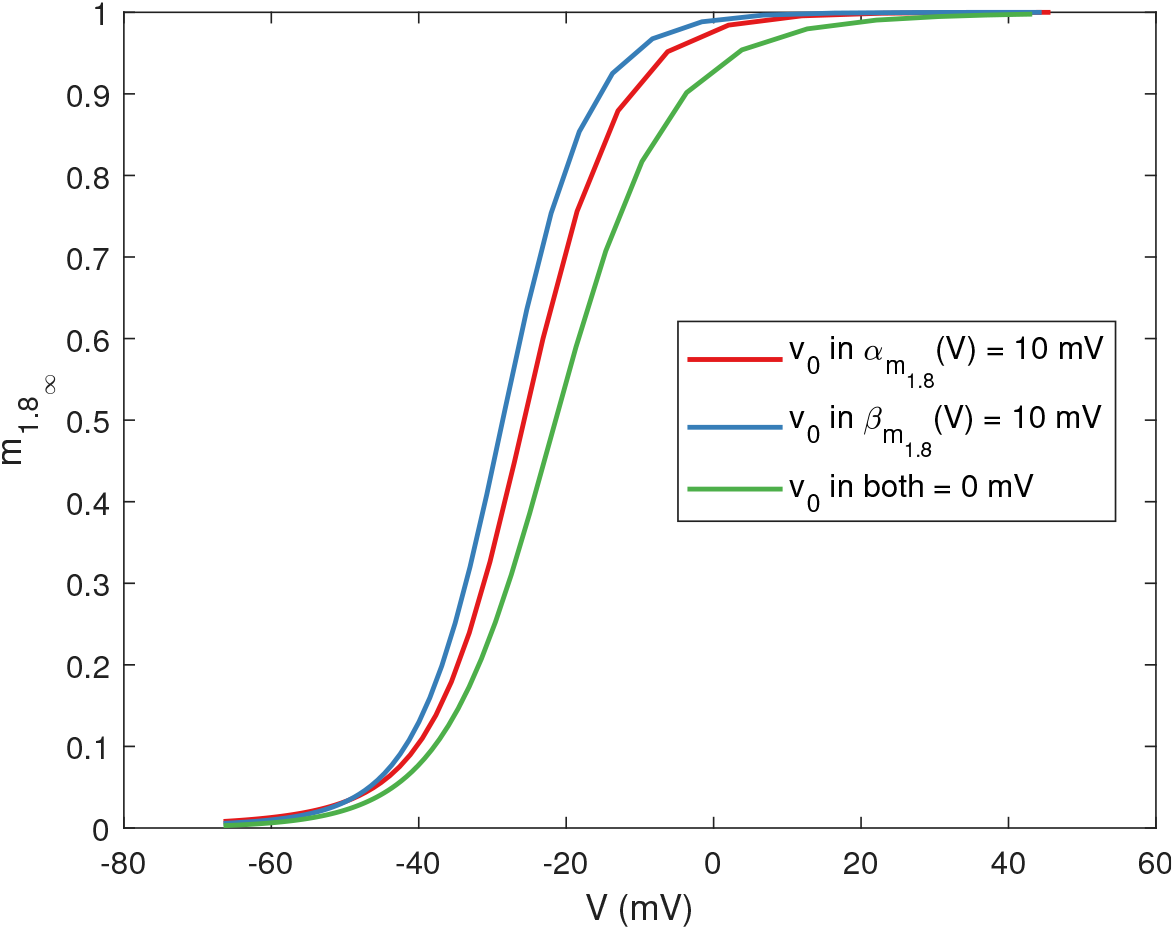
There is a shift in *m*_18_∞__ as a result of an increase in *v*_0_.

The above channel mutation analyses demonstrated the use of bifurcation theory. In particular, two-parameter continuation of bifurcation points identified specific parameters that can shift the bifurcation points and impact the excitability of this neuron.

## 5 Influence of Ion Equilibrium Potentials at *I_ext_* = 0

In this section, we will present the results with *E_Na_* and *E_K_* as the bifurcation parameters and keeping *I_ext_* = 0. These equilibrium potentials are functions of intracellular and extracellular ionic concentrations [33, 17]. A nerve injury can change the ion concentration, subsequently impacting the equilibrium ion potential. A computational study on the nerve injury has been done with Hodgkin-Huxley type equations [58], where alterations in ion concentrations changed the excitability of the neuron. Here, we will demonstrate how sodium and potassium equilibrium ion potentials can potentially alter the pain sensation threshold, using bifurcation theory.

The partial bifurcation diagrams are shown in Fig. 14. The structures are similar to the previous bifurcation diagram in Fig. 9. The diagram consists of a HB with unstable periodic solutions arising from it. There are two LP leading to hysteresis in the unstable steady state solution branch. The unstable periodic solution branch emanating from the HB point ends at an unstable steady state branch, with a turning point (CLP) in between. This end point represents a homoclinic orbit, which is a specific type of periodic solution with infinite period. Another CLP separates unstable and stable periodic solutions. MMO are observed between the HB point and this CLP. In both the cases, an increase in the equilibrium potential leads to generation of periodic firing even without any external stimulus. This is called spontaneous firing, and is indicative of neuropathic pain. In case of neuropathic pain, one may experience a tingling and burning sensation even without any external stimulus. We are not aware of any experimental or computational analysis of equilibrium ion potentials and their relation to pain, although this has been studied theoretically for nerve injury which can lead to pain [58]. The above results suggest a direct link between equilibrium sodium and potassium potential and the periodic firing threshold of a pain-sensing neuron. Further computational and experimental investigation of these results can provide strategies for alleviating neuropathic pain.

**Figure 14:**
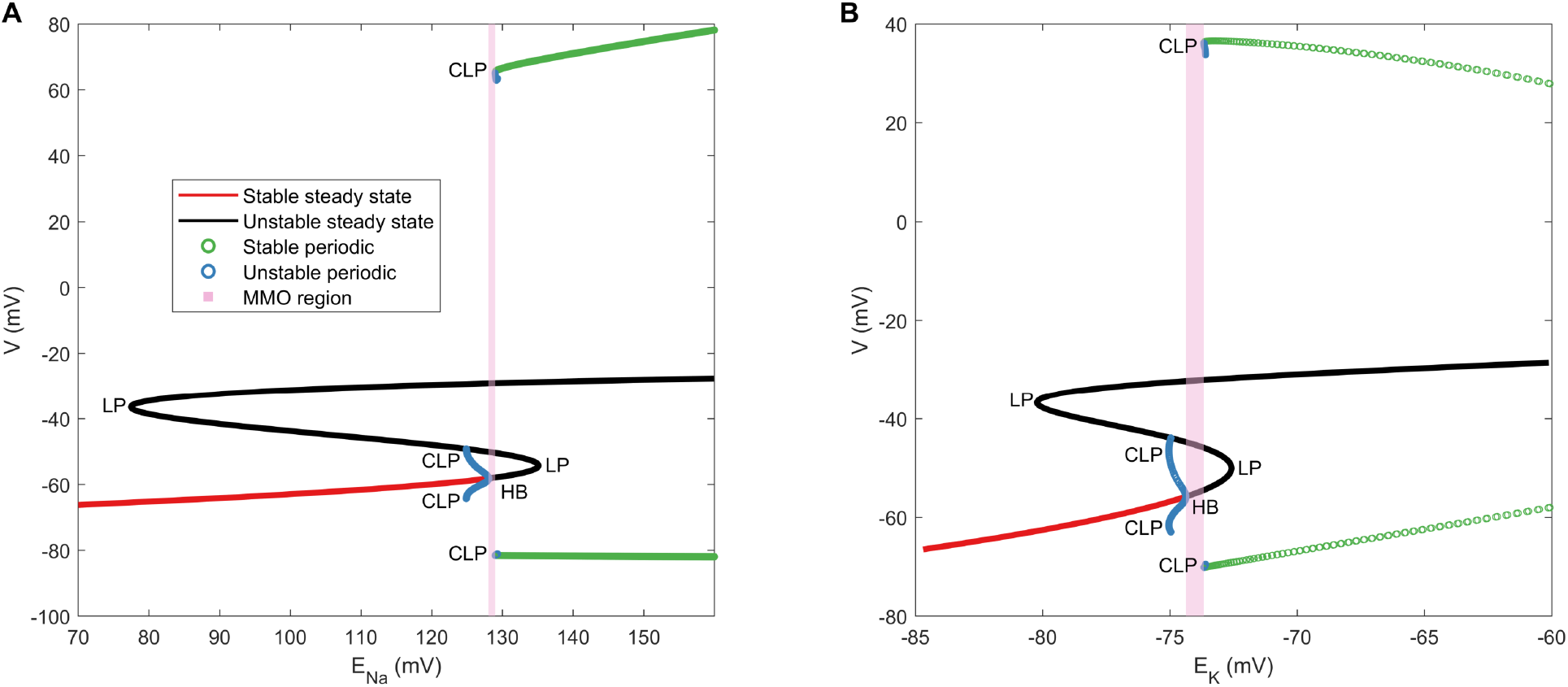
Bifurcation diagram with A: *E_Na_* (HB at *E_Na_* = 128.04 mV and CLP of stable periodic branch at *E_Na_* = 128.96 mV) and B: *E_K_* (HB at *E_K_* = −74.37 mV and CLP of stable periodic branch at *E_K_* = −73.66 mV) as primary bifurcation parameters. HB: Subcritical Hopf bifurcation point, CLP: Cyclic limit point

## 6 Discussions

Pain is an unavoidable aspect of the average human experience. Any form of injury can lead to localized pain. This is the body’s mechanism to begin healing the injured area. However, the pain threshold varies among individuals. The same intensity of a noxious stimulus may not be painful for one individual but may be extremely painful for another. In this paper, we presented a computational approach to understand pain sensation and to detect possible parameters that can shift the pain sensation threshold. To this end, we used bifurcation theory to understand the bifurcations arising in a mathematical model representative of a small DRG neuron’s (pain-sensing neuron) dynamics. The bifurcations of this model can establish the boundary between the sensation of pain and no pain; a stable steady state solution is indicative of no pain, while a periodic solution indicates pain. The frequency of the large amplitude periodic firing further captures the degree of pain. Using this theory, we were also able to identify potential mutation points in sodium channels that can alter the pain sensation threshold. We found evidence in literature for Na_v_1.7 and Na_v_1.8 mutations which were associated with conditions such as chronic pain associated with a burning sensation. We also found that an increase in sodium or potassium equilibrium ion potential can lead to spontaneous firing, an indicator of neuropathic pain. Identifying such sensitive parameters can aid in developing therapeutic drugs that can control the pain sensation threshold.

Our bifurcation analysis has some limitations based on its assumptions. The model used for this study is relatively simple from a physiological point of view. Several other ion channels in addition to the four channels addressed in this paper are present in a small DRG neuron as well. For example, Na_v_1.9, also plays a role in the pain sensation [15, 10, 3] and is present in small DRG neurons but was not included in this model. Furthermore, neuron activity is not only a function of electrochemical reactions but also of biochemical reactions that occur within a neuron and at the synapse, where biochemicals in the form of neurotransmitters participate in synaptic transmission. Additionally, a mutation can also occur at the central nervous system instead of the periphery. In this case, analyzing the synaptic ending can become essential. Since these neurons are relatively long compared to any other cell, starting from the periphery and reaching the spinal cord, organelles are actively transported from one end of the axon to the other. This is called axonal transport. This transport can also get disrupted due to any injury, genetic disorder, or disease, which may lead to neuropathic pain [41]. Lastly, biochemical processes such as release of cytokines and other inflammatory molecules are also involved in generation of inflammatory pain. All the aforementioned mechanisms are potential areas that can benefit from computational modeling and bifurcation theory to find sensitive parameters that impact the pain sensation threshold.

The two parameter approach presented here was useful in identifying potential alterations that can occur in the fast activation of Na_v_1.7 and Na_v_1.8 and can shift the bifurcation points, consequently impacting the pain sensation threshold, which have been observed experimentally in the literature. Our approach’s limitation is in the number of parameters that can be varied simultaneously. Most numerical tools are designed for one and two parameter continuations corresponding to codimension 0 or 1 bifurcation points. For the continuation of higher codimensions, see Krasnyk et al. [35] and references therein.

This work demonstrates the use of bifurcation theory in understanding the variation associated with pain: a vital physiological phenomenon. We identified that alterations in specific model parameters can shift the bifurcation points, which can consequently impact the excitability of a pain-sensing neuron. A thorough analysis using the forgoing approach outlined in this paper corroborated with experimental studies can advance our understanding of the mechanism of pain sensation.

## Supporting information

Supplemental doc

## Acknowledgement

The authors thank Max Planck Institute for Dynamics of Complex Technical Systems, Germany, for sponsoring visits in order to foster discussions and strengthen collaboration on this work, Dr. Haroon Anwar, New Jersey Institute of Technology, USA, for providing directions on model selection, Muriel Eaton and Dr. Yang Yang, Purdue University, USA, for useful discussions on pain sensation and DRG neurons.

XPPAUT[16] was used for numerical integration and bifurcation analysis, MATCONT[12] for performing two-parameter continuation of bifurcation points, and MATLAB[37] for generating the plots. Biological figures were developed using Motifolio (https://www.motifolio.com/) and somersault18:24 (https://www.somersault1824.com/).

## Supplementary Information

XPPAUT and MATCONT codes are available upon request. Model equations and parameter values are mentioned in the supplementary document.

