## Supplemental doc for "Using Bifurcation Theory for Exploring Pain"

### 1 XPPAUT settings

NTST = 100, Method = Stiff, Tolerance = 1e-07, EPSL, EPSU, EPSS = 1e-07, ITMX, ITNW = 20, Dsmin = 1e-05, Dsmax = 0.05. Other settings were same as default. In some cases, there may be a need to adjust Dsmin and Dsmax.

Settings were same for MATCONT, but the equations were re-scaled as shown in Sec. 3.

### 2 Model equations

As described in the main text, the equation for voltage can be written as:

$$c\frac{\mathrm{d}V}{\mathrm{d}t} = \frac{I_{ext}(t)}{A} - (i_{1.7} + i_{1.8} + i_K + i_{KA} + i_l), \tag{1}$$

where, $A$ is the membrane surface area, $t$ is time and $c$ is the specific capacitance. $c\frac{\mathrm{d}V}{\mathrm{d}t}$ is the specific capacitive current, $\frac{I_{ext}(t)}{A}$ is the external current per surface area, and $i_{1.7} + i_{1.8} + i_K + i_{KA} + i_l$ is the ionic current per surface area. The parameter values for this equation are mentioned in Table1. The model equations were obtained from literature[1–3].

The channel specific ionic currents are defined as following:

1. $i_{1.7} = \overline{g}_{1.7} m_{1.7}^3 h_{1.7} s_{1.7} (V - E_{Na})$
2. $i_{1.8} = \overline{g}_{1.8} m_{1.8} h_{1.8} (V - E_{Na})$
3. $i_K = \overline{g}_K n_K (V - E_K)$
4. $i_{KA} = \overline{g}_{KA} n_{KA} h_{KA} (V - E_K)$
5. $i_l = \overline{g}_l (V - E_l)$

Table 1: Voltage dynamics equation parameter values

| Parameter | Value | Units |
|---|---|---|
| $A$ (area) | 2168.00 | $\mu$m$^2$ |
| $C$ | 20.20 | pF |
| $E_{Na}$ | 67.10 | mV |
| $E_K$ | -84.70 | mV |
| $E_l$ | -58.91 | mV |
| $\overline{g}_{1.7}$ | 18.00 | mS/cm$^2$ |
| $\overline{g}_{1.8}$ | 7.00 | mS/cm$^2$ |
| $\overline{g}_K$ | 4.78 | mS/cm$^2$ |
| $\overline{g}_{KA}$ | 8.33 | mS/cm$^2$ |
| $\overline{g}_l$ | 0.0575 | mS/cm$^2$ |

Table 2: Gating variables parameters

| Parameter | $k_1$ ($\text{ms}^{-1}$) | $k_2$ ($\text{ms}^{-1}$) | $k_3$ (mV) | $k_4$ (mV) |
|---|---|---|---|---|
| $\alpha_{m_{1.7}}$ | 0 | 15.5 | -5 | -12.08 |
| $\beta_{m_{1.7}}$ | 0 | 35.2 | 72.7 | 16.7 |
| $\alpha_{h_{1.7}}$ | 0 | 0.38685 | 122.35 | 15.29 |
| $\beta_{h_{1.7}}$ | -0.00283 | 2.00283 | 5.5266 | -12.70195 |
| $\alpha_{s_{1.7}}$ | 0.00003 | 0.00092 | 93.9 | 16.6 |
| $\beta_{s_{1.7}}$ | 132.05 | -132.05 | -384.9 | 28.5 |
| $\alpha_{m_{1.8}}$ | 2.85 | -2.839 | -1.159 | 13.95 |
| $\beta_{m_{1.8}}$ | 0 | 7.6205 | 46.463 | 8.8289 |

The final equation for voltage is the following:

$$\begin{aligned} c\frac{\mathrm{d}V}{\mathrm{d}t} = \frac{I_{ext}(t)}{A} &- (\overline{g}_{1.7}m_{1.7}^3h_{1.7}s_{1.7}(V - E_{Na}) + \overline{g}_{1.8}m_{1.8}h_{1.8}(V - E_{Na}) \\ &+ \overline{g}_K n_K(V - E_K) + \overline{g}_{KA}n_{KA}h_{KA}(V - E_K) \\ &+ \overline{g}_l(V - E_l)) \end{aligned}$$

For any gating variable $x$ ($x = m_{1.7}, h_{1.7}, s_{1.7}, m_{1.8}, h_{1.8}, n_K, n_{KA}, h_{KA}$), the equation for gating variable dynamics can be written as:

$$\frac{\mathrm{d}x}{\mathrm{d}t} = \frac{x_\infty(V) - x}{\tau_x(V)}, \tag{2}$$

where

$$x_\infty(V) = \frac{\alpha_x(V)}{\alpha_x(V) + \beta_x(V)}, \tag{3}$$

and

$$\tau_x(V) = \frac{1}{\alpha_x(V) + \beta_x(V)} \tag{4}$$

The general form of $\alpha_x(V)$ and $\beta_x(V)$ is the following:

$$k_1 + \frac{k_2}{1 + \exp\frac{V + k_3}{k_4}}, \tag{5}$$

where, $k_1$, $k_2$, $k_3$, $k_4$ are constants. Most of the variables follow the above form, however, there are some exceptions, which we will mention in the following subsections. For the rest, the parameter values are mentioned in Table 2.

### 2.1 Nav1.8 kinetics

$$\tau_{h_{1.8}}(V) = 1.218 + 42.043 \exp\left(-\frac{(V + 38.1)^2}{2 \cdot 15.19^2}\right) \tag{6}$$

$$h_{1.8_\infty}(V) = \frac{1}{1 + \exp\left(\frac{V + 32.2}{4}\right)} \tag{7}$$

### 2.2 K kinetics

$$\alpha_{n_K}(V) = \frac{0.001265\,(V + 14.273)}{1 - \exp\left(-\frac{V + 14.273}{10}\right)} \tag{8}$$

If $\alpha_{n_K}(V) = -14.273$, $\alpha_{n_K}(V) = 0.001265 \times 10$

$$\beta_{n_K}(V) = 0.125 \exp\left(-\frac{V+55}{2.5}\right) \tag{9}$$

$$n_{K_\infty}(V) = \frac{1}{1+\exp\left(-\frac{V+14.62}{18.38}\right)} \tag{10}$$

$$\tau_{n_K}(V) = \frac{1}{\alpha_{n_K}+\beta_{n_K}} + 1 \tag{11}$$

#### 2.3 KA kinetics

$$n_{KA_\infty}(V) = \left(\frac{1}{1+\exp\left(-\frac{V+5.4}{16.4}\right)}\right)^4 \tag{12}$$

$$\tau_{n_{KA}}(V) = 0.25 + 10.04 \exp\left(-\frac{(V+24.67)^2}{2\cdot 34.8^2}\right) \tag{13}$$

$$h_{KA_\infty}(V) = \frac{1}{1+\exp\left(\frac{V+49.9}{4.6}\right)} \tag{14}$$

$$\tau_{h_{KA}}(V) = 20 + 50 \exp\left(-\frac{(V+40)^2}{2\cdot 40^2}\right) \tag{15}$$

If $\tau_{h_{KA}}(V) < 5, \tau_{h_{KA}}(V) = 5$.

### 3 Non-dimensional equations

To non-dimensionalize, let us introduce some constants $k_v$, $k_t$, $g$, $T_x$ ($x = m_{1.7}, h_{1.7}, s_{1.7}, m_{1.8}, h_{1.8}, n_K, n_{KA}, h_{KA}$). The non-dimensional variables will be: $V = k_v \cdot v$, $E_X = k_v \cdot \tilde{E}_X$, ($X = Na, K, l$), $t = k_t \cdot \tilde{t}$, $\overline{g}_y = g \cdot \tilde{g}_y$ ($y = 1.8, 1.7, K, KA, l$), $I_{ext} = \tilde{I}_{ext}/(k_v \cdot g \cdot A)$, $\tau_x = T_x \cdot \tilde{\tau}_x$. Then, the equations will be:

$$\begin{aligned}\frac{\mathrm{d}v}{\mathrm{d}\tilde{t}} = \frac{k_t \cdot g}{c}\big[\tilde{I}_{ext} - (&\tilde{g}_{1.7} m_{1.7}^3 h_{1.7} s_{1.7}(v-\tilde{E}_{Na}) + \tilde{g}_{1.8} m_{1.8} h_{1.8}(v-\tilde{E}_{Na}) \\ &+ \tilde{g}_K n_K(v-\tilde{E}_K) + \tilde{g}_{KA} n_{KA} h_{KA}(v-\tilde{E}_K) \\ &+ \tilde{g}_l(v-\tilde{E}_l))\big]\end{aligned} \tag{16}$$

$$\frac{\mathrm{d}m_{1.7}}{\mathrm{d}\tilde{t}} = \frac{k_t}{T_{m_{1.7}}}\frac{(m_{1.7_\infty}(v) - m_{1.7})}{\tilde{\tau}_{m_{1.7}}(v)} \tag{17}$$

$$\frac{\mathrm{d}h_{1.7}}{\mathrm{d}\tilde{t}} = \frac{k_t}{T_{h_{1.7}}}\frac{(h_{1.7_\infty}(v) - h_{1.7})}{\tilde{\tau}_{h_{1.7}}(v)} \tag{18}$$

$$\frac{\mathrm{d}s_{1.7}}{\mathrm{d}\tilde{t}} = \frac{k_t}{T_{s_{1.7}}}\frac{(s_{1.7_\infty}(v) - s_{1.7})}{\tilde{\tau}_{s_{1.7}}(v)} \tag{19}$$

$$\frac{\mathrm{d}m_{1.8}}{\mathrm{d}\tilde{t}} = \frac{k_t}{T_{m_{1.8}}}\frac{(m_{1.8_\infty}(v) - m_{1.8})}{\tilde{\tau}_{m_{1.8}}(v)} \tag{20}$$

$$\frac{\mathrm{d}h_{1.8}}{\mathrm{d}\tilde{t}} = \frac{k_t}{T_{h_{1.8}}}\frac{(h_{1.8_\infty}(v) - h_{1.8})}{\tilde{\tau}_{h_{1.8}}(v)} \tag{21}$$

$$\frac{\mathrm{d}n_K}{\mathrm{d}\tilde{t}} = \frac{k_t}{T_{n_K}} \frac{(n_{K_\infty}(v) - n_K)}{\tilde{\tau}_{n_K}(v)} \tag{22}$$

$$\frac{\mathrm{d}n_{KA}}{\mathrm{d}\tilde{t}} = \frac{k_t}{T_{n_{KA}}} \frac{(n_{KA_\infty}(v) - n_{KA})}{\tilde{\tau}_{n_{KA}}(v)} \tag{23}$$

$$\frac{\mathrm{d}h_{KA}}{\mathrm{d}\tilde{t}} = \frac{k_t}{T_{h_{KA}}} \frac{(h_{KA_\infty}(v) - h_{KA})}{\tilde{\tau}_{h_{KA}}(v)} \tag{24}$$

$k_v$ is set to 150 mV (approximately the difference between $E_{Na}$ and $E_K$), $k_t$ as 30 ms (approximately the time covered by one action potential) and $g$ as the $\overline{g}_{1.7} = 18$ mS/cm$^2$ (largest value of maximal conductance). Approximate values of the remaining constants can be set as: $T_{m_{1.7}} = 0.2$, $T_{h_{1.7}} = 40$, $T_{s_{1.7}} = 9000$, $T_{m_{1.8}} = 1$, $T_{h_{1.8}} = 40$, $T_{n_K} = 300$, $T_{n_{KA}} = 10$, $T_{h_{KA}} = 70$ ms. The approximate time scale of evolution of the variables $v, m_{1.7}, h_{1.7}, s_{1.7}, m_{1.8}, h_{1.8}, n_K, n_{KA}, h_{KA}$ can be calculated from the constants on the right hand side of each of these equations, which are equal to 580, 150, 0.75, 0.0033, 30, 0.75, 0.1, 3, 0.42 respectively. The order of magnitude provides insight into the speed of each of these variables. $v$ and $m_{1.7}$ are the fastest and $s_{1.7}$ the slowest.
